# Unlocking the genomic taxonomy of the *Prochlorococcus* collective

**DOI:** 10.1101/2020.03.09.980698

**Authors:** Diogo Tschoeke, Livia Vidal, Mariana Campeão, Vinícius W. Salazar, Jean Swings, Fabiano Thompson, Cristiane Thompson

**Affiliations:** Laboratory of Microbiology. SAGE-COPPE and Institute of Biology. Federal University of Rio de Janeiro. Rio de Janeiro. Brazil. Av. Carlos Chagas Fo 373, CEP 21941-902, RJ, Brazil; Laboratory of Microbiology, Ghent University, Gent, Belgium

**Author notes:** These authors contributed equally (Diogo Tschoeke, Livia Vidal).

**Keywords:** *Prochlorococcus* collective, ecogenomics, genomic taxonomy, evolution

## Abstract

*Prochlorococcus* is the most abundant photosynthetic prokaryote on our planet. The extensive ecological literature on the *Prochlorococcus* collective (PC) is based on the assumption that it comprises one single genus comprising the species *Prochlorococcus marinus*, containing itself a collective of ecotypes. Ecologists adopt the distributed genome hypothesis of an open pan-genome to explain the observed genomic diversity and evolution patterns of the ecotypes within PC. Novel genomic data for the PC prompted us to revisit this group, applying the current methods used in genomic taxonomy. As a result, we were able to distinguish the five genera: *Prochlorococcus, Eurycolium, Prolificoccus, Thaumococcus* and *Riococcus*. The novel genera have distinct genomic and ecological attributes.

## INTRODUCTION

*Prochlorococcus* cells are the smallest cells that emit red chlorophyll fluorescence, which is how they were discovered approximately 30 years ago [1, 2]. It is the smallest and most abundant photosynthetic microbe on Earth. The oxytrophic free-living marine *Prochlorococcus marinus* contain divinyl chlorophyll a (chl a2) and both monovinyl and divinyl chlorophyll b (chl b2) as their major photosynthetic pigments rather than chlorophyll a and phycobiliproteins, which are typical of cyanobacteria, allowing them to adapt to different light intensities [1, 3–6], particularly to capture the blue wavelengths prevailing in oligotrophic waters [7]. *Prochlorococcus* strains have been cultured and are available in culture collections, but the limited number of cultures does not reflect the true phenotypic and genomic diversity found in the ocean [8]. *Prochlorococcus* is a smaller, differently pigmented variant of marine *Synechococcus* [2]. The pioneering work on the eco-genomics of *Prochlorococcus* was performed by Chisholm’s group at MIT [2, 9–14]. “When I look at our emerald green laboratory cultures of captive *Prochlorococcus*, my mind quickly turns to their wild cousins, drifting freely in the world oceans. I am reminded that while they carry out a respectable fraction of the photosynthesis on our planet, they escaped our attention” [15]. The taxonomy of the PC has been neglected in the last three decades. The designation PC was first coined by Kashtan et al. [11]. This collective comprises the genera, *Prochlorococcus, Prolificoccus, Eurycolium*, and *Thaumococcus* [16–18], a group of picocyanobacteria that radiated from *Synechococcus* millions of years ago. We preferer to use the neutral term “collective” here instead of “community”, “complex” or any other term.

### “Speciation” within PC

Several authors have commented on speciation within this group of picocyaonobacteria. Berube et al. [19] argue that speciation is likely driven by a combination of vertical inheritance and gene loss and is rarely due to horizontal gene acquisition driven by non-homologous recombination. Speciation and niche partitioning involve brief periods of enhanced drift and (rapid) change that have facilitated, e.g., the genomic rearrangement of nitrate assimilation genes. Thompson et al. [20] demonstrated that *Prochlorococcus* from the Red Sea is not genetically isolated, despite its endemic genomic adaptations to the unique combination of high solar irradiance, high temperature, high salinity, and low nutrient levels in this sea. The authors consider orthologous groups of genes found in Red Sea *Prochlorococcus* to be cosmopolitan. Species exist as clusters with genetic and ecological similarity, and speciation seems mainly to be driven by natural selection, regardless of the balance between horizontal and vertical descent. “If complete boundaries to gene flow take some time to emerge, we can think of gene sets rather than whole genomes as the units that inhabit ecological niches. If the gene flow boundaries never emerge, speciation does not occur (i.e. we are left with one species, not two) and this corresponds to the gene ecology model” of Shapiro et al. or to Cohan’s stable ecotype model of speciation [21–24]. Shapiro [25] further developed the ideas that “no genome is an island” and that natural genetic engineering responds to the ecological challenges that arise in the life histories of all evolving organisms.

Speciation within the PC may be influenced by co-evolution with *Pelagibacter*, a member of the SAR 11 clade [26–28]. The role of a cryptic nitrogen cycle has been elucidated recently [3]: *Pelagibacter* mainly uses nucleotide (purine) catabolism to obtain reducing power and releases urea, while *Prochlorococcus* utilizes this urea. The authors postulate the existence of a cryptic nitrogen cycle that transfers reducing power from *Prochlorococcus* to *Pelagibacter* and returns nitrogen to *Prochlorococcus*. This positive feedback loop would strengthen the co-evolution between these two abundant oceanic microbes.

Which forces have shaped the massive cell size/genome reductions and gigantic free-living populations of the PC during its millions years of evolution? The streamlining theory is generally cited to explain the massive genomic DNA losses and cell size reduction and the accompanying AT increase [29]. Low genetic drift and strong purifying selection have been postulated. The major driving force for genome reduction seems to be the substantial economy in energy and materials that has occurred, as the AT increase economizes nitrogen [30].

The PC is cited as an example of the evolution of dependencies of free-living organisms through adaptive gene losses, better known as the Black Queen hypothesis [31, 32], summarized by Shapiro (2019) [25] as “no genome is an island”, to stress continuous biosphere communications during evolution.

Most literature on PC is based on the assumption that it comprises a single genus, in which the species *Prochlorococcus marinus* is within a “collective” of ecotypes. The main reason for this concept is that 16S rRNA gene sequence similarities within this group are >97%. To describe the observed diversity in the PC, ecologists adopt the distributed genome hypothesis of an open bacterial pan-genome (supra-genome) [12, 14]. In this hypothesis, the pan-genome is the full complement of genes that each member of the population contributes to and draws genes from. No single isolate contains the full complement of genes, which contributes to high diversity and provides selective advantages [33]. Described in its extreme form, the pan genome of the bacterial domain is of infinite size, and the bacteria as a whole present an open pan-genome (supragenome), in which the constant “rain” of genetic material in genomes from a cloud of frequently transferred genes enhances the chance of survival of a species [34]. For scientific progress, the use of these metaphors is necessary to clarify ideas and formulate new hypotheses. The PC case illustrates the continuously growing gap between prokaryotic taxonomy as it is practised today and in-depth eco-genomic studies. Genomic clusters are referred to as ecotypes, ecospecies, sub-ecotypes, clades, populations, subpopulations, OTUs or CTUs. This informal nomenclature finds its way into databases, adding further nomenclatural chaos to “terrae incognitae” [35, 36] comprising these picocyanobacteria.

In their theory of self-amplification and self-organization of the biosphere, Braakman et al. [3], hypothesize that *Prochlorococcus* clades living in surficial oceanic waters adapted to high light and low nutrient levels have evolved from more ancient *Prochlorococcus s*ubpopulations (∼800 million years ago) inhabiting deeper oceanic areas, with lower light and higher nutrient concentrations. They refer to these evolutionary paths as “niche constructing adaptive radiations”. Sun & Blanchard [37] hypothesise three phases in the evolutionary history of the PC: genome reduction, genome diversification and ecological niche specialization, all due to different evolutionary forces. The PC has adapted to live under different light intensities, sea water temperatures and nutrient levels, and the abundance of the different ecotypes varies markedly along gradients of these parameters [38–44]. However, only high light (HL)-adapted and low light (LL)-adapted ecotypes were initially distinguished [45]. At present, twelve phylogenetic ecotypes are delineated: six high light (HL)-adapted ecotypes and six low light (LL)-adapted ecotypes [46]. These ecotypes are based on the phylogeny of the 16S/23S rRNA internal transcribed spacer region, differences in behaviour towards light, whole-genome analysis, and physiological properties, combined with their environmental distribution [42, 44, 47]. Not only light adaptation but also temperature adaptation has been recognized in the two dominant near-surface HL ecotypes, the HL I ecotype *Eurycolium* eMED4 and the HL II ecotype *Eurycolium* eMIT9312, which both exhibit growth maxima at approximately 25 °C, but *Eurycolium* eMED4 appears to be adapted to thrive at much lower temperatures (approx. 14 °C), whereas *Eurycolium* eMIT9312 is able to grow at 30 °C [48]. These ecotypes appear to have nearly identical genomes [9], in which the major differences are restricted to five genomic islands, but show marked phenotypic differences.

The relative abundance of cells belonging to the different ecotypes has been estimated in several oceanic regions, revealing patterns that agree for the most part with the HL/LL phenotypes. HL-adapted cells dominate the surface mixed layer, and LL-adapted cells most often dominate deeper waters [46, 49–52]. By combining the physiological features of isolates and ecotype abundance studies in the ocean, it was shown that temperature, in addition to light, is an important determinant of the abundance of these ecotypes in oceans [53, 54]. There is also tremendous variability within ecotypes, particularly with regard to nutrient acquisition [55], susceptibility to predation or phage infection [56], and modes of interaction with other members of the community [57], including interactions with *Alteromonas* for protection against ROS [58, 59]. *Prochlorococcus* and *Synechococcus* can use different organic compounds containing key elements to survive in oligotrophic oceans, and can also take up glucose and use it as a source of carbon and energy[60].

Antisense RNA in *Prochlorococcus* is said to protect a set of host mRNAs and viral transcripts from RNAse degradation during phage infection [61]. In addition, CRISPR-Cas (clustered regularly interspaced short palindromic repeats, CRISPR-associated genes), an extremely adaptable defense system, is widely distributed in cyanobacteria [62]. The CRISPR-Cas system is found in the majority of cyanobacteria except for *Synechococcus* and *Prochlorococcus*, in which cyanophages are a known force shaping evolution [62]. However, more recently, CRISP-Cas systems have been found in novel *Synechococcus* genomes, indicating the possibility of additional mechanisms for genome plasticity [63].

Kashtan et al. [11] studied the genomic diversity of single *Prochlorococcus* cells in samples collected in the Bermuda-Atlantic Time-Series Study (BATS) site in summer, spring, and winter, and hundreds of new *Prochlorococcus* “subpopulations” with distinct genomic backbones were found. These subpopulations are estimated to have diverged millions of years ago, suggesting ancient, stable niche partitioning followed by selection (not by genetic drift). The authors suggested that the global PC may be conceived as an assortment of thousands of species. They argued that such a large set of coexisting species with distinct genomic backbones is a characteristic feature of free-living prokaryotes with very large population sizes living in highly mixed habitats. In another study by these authors focused on the Pacific Ocean, they concluded that the Atlantic and Pacific Oceans are occupied by different PC populations that experienced separate evolutionary paths for a few million years [13].

Phylogeography seems to be echoed in the extensive genome diversity of the PC identified in 226 metagenomic samples from the GOS (Global Ocean Sampling) expedition [44]. In another paper, Kent et al. [64] studied the co-occurrence of *Prochlorococcus* and *Synechococcus* in 339 samples from three cruises covering equatorial, subtropical gyre, and colder nutrient-rich mid-latitude regions in the Atlantic and Pacific using the phylogenetic marker gene *rpoC*1, a single-copy gene encoding the γ subunit of RNA polymerase. They hypothesized that where *Prochlorococcus* and *Synechococcus* co-occur, parallel evolutionary diversification occurs, caused by the same stressors. The authors distinguish three major matching surface ecotypes: *Prochlorococcus* HLIII/IV (corresponding to *Synechococcus* CRD1); HLII (*Synechococcus* clade II/III) and HLI (*Synechococcus* I/IV). This parallel phylogeography extends from the ecotype level further into finer genomic networked diversity.

### Principles of genomic prokaryote taxonomy

From 1970 to the present, the need to classify, name and identify prokaryotes has been based on a pragmatic polyphasic consensus approach [65]. This non-theory-based approach aims to integrate phenotypic, genotypic, and phylogenetic and ecological characteristics to establish stable and informative classification systems. The central unit is the prokaryotic species. In this procedure, the definition of prokaryotic species is based mainly on DNA–DNA hybridization (DDH) and 16S rRNA sequence similarity: a species is defined as a monophyletic group of strains sharing high phenotypic and genomic similarity (>70% DDH, >98.3% 16S rRNA sequence similarity) [66–70].

Prokaryote taxonomy is becoming more genome based, considering the genomes themselves as the ultimate end products of microbial evolution [68, 71, 72]. Such an approach offers the opportunity to obtain a more natural taxonomic system and allows the definition of consistent and stable monophyletic species and genera with maximal utility for its users (see 47, 48, 78–98). Cohan has extensively developed the ecotype or ecospecies theory for prokaryotes, in which “ecotypes are the most newly divergent populations that are distinct ecologically from one another [21, 73]. In this theory, the splitting of one ecotype into two is the fundamental diversity-creating process of speciation in bacteria. In his stable ecotype model, ecotypes (ecospecies) are long lived, and different ecotypes coexist indefinitely. They are cohesive by virtue of periodic selection events that purge the ecotype of sequence diversity. In this case, the ecotype has all the characteristics of a species [23, 73]. Ecotypes have been discerned among the PC collective.

### The place of the PC in marine microbial taxonomy

The main motivation for the present study arose from the consideration that the taxonomy of *Prochlorococcus* has remained relatively neglected and unresolved compared to the tremendous advances in marine microbial taxonomy achieved for other groups, e.g., *Alteromonas* (N=38 species) [74], *Colwellia* (N=21 species) [75], *Endozoicomonas* (N=9 species) [76], *Marinobacter* (N=45 species) [77], *Marinobacterium* (N=17 species) [78], *Marinomonas* (N=30 species) [79], *Pseudoalteromonas* (N=53 species) [80], *Pseudovibrio* (N=6 species) [81], *Roseobacter* (N=5 species) [82], *Ruegeria* (N=22 species) [83], *Shewanella* (N=67 species) [84], *Salinispora* (N=3 species) [85], *Streptomyces* (N=848 species) [86], *Thalassomonas* (N=9 species) [87], and the vibrio clade (N=199 species, including the genera *Vibrio, Photobacterium, Enterovibrio, Aliivibrio, Thaumasiovibrio*, and *Grimontia*) [88]. Even marine prokaryotes living in extreme oceanic environments such as the OMZ, sulfate-reducing bacteria, N cycle bacteria, and deep sea hydrothermal vent sulfur-oxidizing *Sulfurovum* (N=4 species) [89] and *Sulfurimonas* (N=4 species) [90] have been the subject of advanced taxonomic classification. Nevertheless, even in the well-studied Gram-negative *Roseobacter* [91] and Gram-positive *Streptomyces* [92] genera, the paraphyletic nature of these genera leads to noncoherent assignment of scientific names and causes confusion when interpreting their ecology or evolutionary biology.

As new habitats and niches are found for *Prochlorococcus*, it is important to establish a solid taxonomic system to contribute to the interpretation of the ecology of the PC. Novel PC genomic data allowed us to address the hypothesis that novel genera and species are present within the PC. In this review, we revisit the current taxonomy of the PC and propose the creation of a new genus to encompass the clearly recognized clusters. The novel genera have distinct genomic and ecological attributes.

## METHODS

### Genome quality assessment

A total of 208 genomes and the corresponding metadata were downloaded from NCBI GenBank [93]. Contigs or scaffold files were input into Prodigal software v2.6.3 [94] for gene and protein prediction. To check quality of the genomes, completeness, contamination and GC%, the genomes were submitted to CheckM v1.0.11 [95] using default parameters for this estimation. CheckM uses a broader set of marker genes specific to the position of a genome within a reference genome tree. The completeness is measured using a set of marker genes that are expected to be present. This expectation can be defined along an evolutionary gradient, ranging from highly conserved genes to species-specific genes. The genomes used are of high quality and have ≥ 85% of completeness, ≥ 1Mb of size and ≤ 5% of contamination.

### Average amino acid identity analyses (AAI)

The average amino acid identity was calculated as described previously for all 208 genomes [96]. All PC genomes were used to calculate the amino acid average identity using the *CompareM* tool (https://github.com/dparks1134/CompareM). The result was used to plot a dendrogram. The *CompareM* output was converted to a dissimilarity square matrix. The *spatial.distance.squareform* function from *scipy* package v1.1.0 was used in order to obtain a condensed list from the lower triangular elements of the matrix. A complete linkage clustering was performed using *cluster.hierarchy.linkage* function from *scipy* with the output of *squareform* and cityblock metric distance. The figure that represents the linkage matrix was plotted with *cluster.hierarchy.dendrogram* function from *scipy*. The intraspecies limit is assumed as ≥95%, whereas genera delimitation is assumed as ≥70%. Genera were delineated only for those lineages exhibiting at least 3 representative genome sequences.

### Phylogenomic reconstruction

The genome sequences of five representative PCs corresponding to the type species of the five genera were selected based on AAI analysis. The MLSA approach was based on the concatenated sequences of four house-keeping genes (*gyrB, pyrH, recA, rpoB*) with intraspecies relatedness ranging from >95 to 100 % MLSA[16]. This was achieved by using tBLASTn [97]. A nucleotide alignment was then made for each of the four genes using MAFFT [98], and these alignments concatenated using MAFFT with default parameters and a custom Perl script. The tree was built from the concatenated alignment using FastTree 2 [99] with default parameters and rooted on *S. elongatus* PCC 6301 using ETE 3 [100]. To build the core genome tree, we used the GToTree package [101] with default parameters. The 208 selected genomes were searched against a Hidden Markov Model of 249 Cyanobacterial marker genes using HMMER3 [102]. A concatenated protein alignment from the 249 genes was constructed using Muscle [103] and subsequently trimmed using TrimAl [104] (Gutierrez et al. 2009). The alignment was then used to construct a tree using FastTree 2 [99] with default parameters and the pairwise distance matrix using MEGA 6.0 [105]. The tree was rooted on S. elongatus PCC 6301 using ETE 3 [100]. Part of the processing was done with GNU Parallel [106].

### *In silico* phenotype

The phenotype was predicted utilizing the sequences of the nitrogen (*narB, nirA, ntcA, glnB, amt, cynS, ureA, glnA, glsF, gdhA*), phosphorous (*phoB, ptrA, phoR*), arsenate (*arsC*) and iron (*piuC, som*) related genes of *Prochlorococcus* obtained from UniProt and GenBank as queries for a local blastp [107] search against all PC ORF-predicted genomes. The results were parsed with Python scripts, and the sequences corresponding to the blastp hits were extracted from the respective genomes. The extracted sequences were used to perform a blastp search against the nr database to confirm the functional annotation. Manual verification of these results was performed. The confirmed results were used to build a matrix of presence/absence for the genes related to the indicated proteins in each PC genome.

## RESULTS AND DISCUSSION

### A proposal of a new taxonogenomic framework for the PC

The availability of *Prochlorococcus* genomes labelled *Prochlorococcus* in public databases prompted us to test the hypothesis that the PC comprises a greater number of stable genera and species than has been previously recognized (Table 1 and S1). We first collected all the publicly available genomes designated as *Prochlorococcus* and applied the theory-free genomic methodologies and principles discussed previously. The *Prochlorococcus* genomes came from samples distributed around the world, especially in the Pacific and Atlantic oceans (Table S1). First, the GC% content of the genomes was determined. A phylogenomic framework was established using genomic signatures, i.e., average amino acid identity (AAI), MLSA and core genome-based tree. We then classified the genomes into different genera and species using relevant AAI and phenotypic information obtained directly from the genomes.

**Table 1.** Proposal of a new PC taxonomic classification. t• New taxonomic classification proposed by Walter et al. (2017). a: Average genome size was calculated using all genomes from each genus. The standard deviation was represented by ± symbol.

| Genus (No. of species) | No. of genomes | Type species | Type strain | Range % mol GC | Average length (Mbp) <sup>a</sup> |
| --- | --- | --- | --- | --- | --- |
| *Eurycolium* (97) | 156 | *Eurycolium pastoris* • | CCMP1986<sup>T</sup> | 30-33 | 1.5±0.1 |
| *Prochlorococcus* (04) | 10 | *Prochlorococcus marinus* | CCMP 1375<sup>T</sup> | 36-37 | 1.7±0.07 |
| *Prolificoccus* (16) | 25 | *Prolificoccus proteus*• | NATL2A<sup>T</sup> | 34-55 | 1.7±0.1 |
| *Thaumococcus* (02) | 14 | *Thaumococcus swingsii*• | MIT 9313<sup>T</sup> | 50-50.7 | 2.5±0.06 |
| *Riococcus* gen. nov. (03) | 03 | *Riococcus ceticus* sp. nov. | MIT9211<sup>T</sup> | 37-38 | 1.6±0.1 |

### GC content reveals great genomic diversity in the PC

One of the most striking findings of our study came from the analysis of GC content, which ranged from 30% to 50.7% (Figure 1 and Table 1). The distribution of the GC values of all the genomes comprised five groups: i. the majority (75% of the genomes) exhibited 30 to 33% GC, comprising the genus *Eurycolium*; ii. a second group (12% of the genomes) exhibited 34 to 35% GC, comprising the genera *Prolificoccus*; iii. a third group *Prochlorococcus* (4.8% of the genomes) presented 36 - 37% GC; iv. a fourth group *Riococcus* (1.4% of the genomes) exhibited 37 to 38% GC; and v. the fifth group (6.7% of the genomes) exhibited 50 to 50.7% GC, which was characteristic of the genera *Thaumococcus*. The average genome sizes of the first four GC groups were similar, whereas the genome sizes of the fifth GC group were almost two-fold larger. This GC analysis provides a clear indication of the taxonomic width of the PC. This diversification is the result of millions of years of evolution. Mutational bias is dominated by GC > AT transitions even in bacteria with a high % GC content [108, 109]. This means that there must be some pressure to maintain high % GC contents in those prokaryotes that exhibit high % GC values. Why this is so remains a mystery, but Hershberg & Petrov noted that selective forces consistently operate both genome-wide and over long periods of time [109]. The nature of such selection remains obscure. If natural selection plays such a strong role in determining % GC content, it suggests that there are no truly neutrally evolving sites.

**Figure 1.**
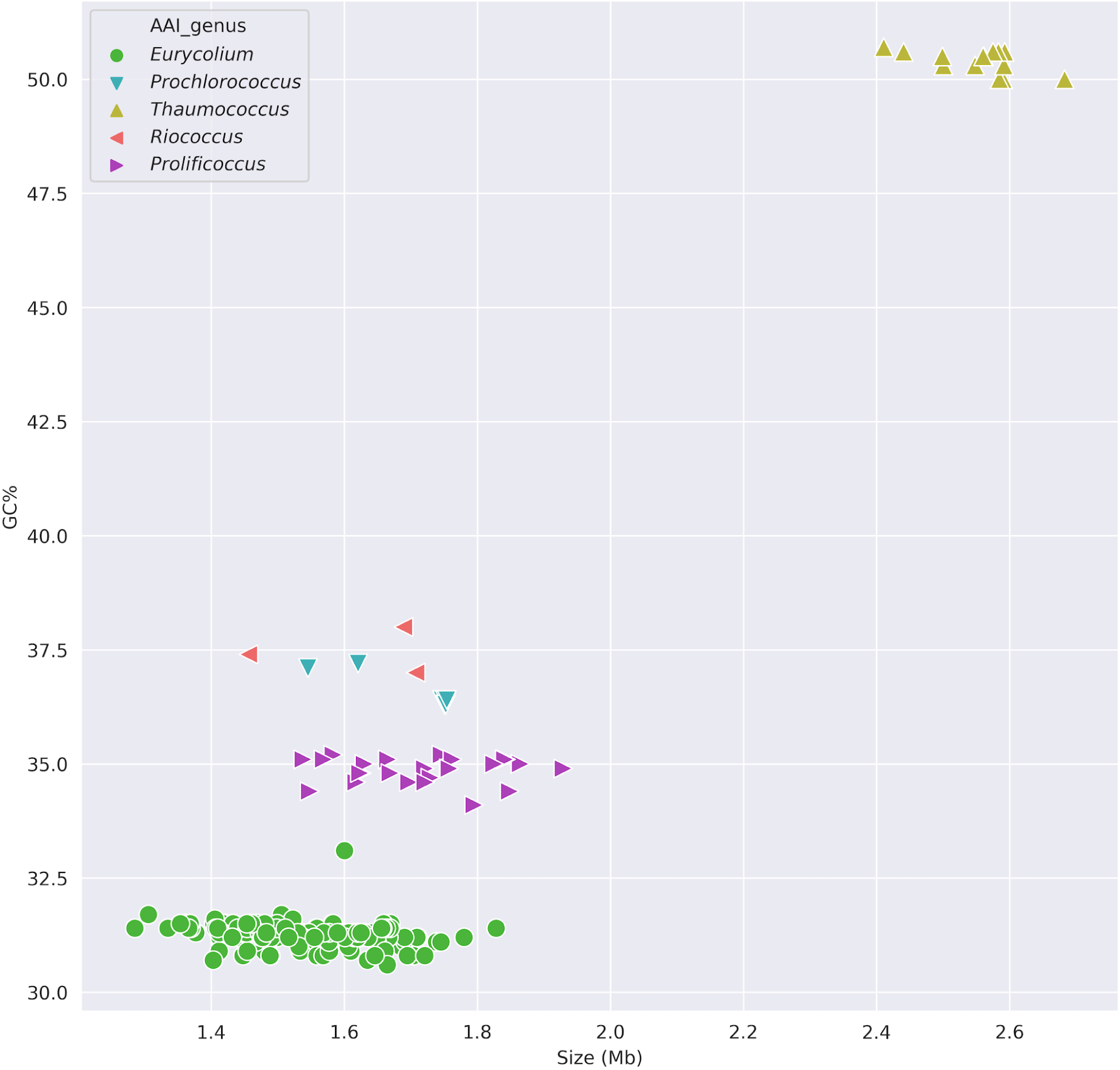
Genome clustering in five groups based on and GC content and genome size. The colors (green, red, blue, purple and yellow) represent the five groups. The five different symbols represent the genera where each genome were classified.

### Average amino acid identity of PC

The species and genus delimitations were based on clusters with three or more genomes that shared ≥95% or ≥70% amino acid average identity (AAI), respectively. Here, we revisit the current taxonomy of the PC and propose the creation of a new genus *Riococcus* and 113 novel species (N=122 species) (Table 1 and S1). We confirmed the previous proposal of Walter et al. [18], and we now expand the classification of PC (Figure 2). *Eurycolium* (N=97) and *Prolificoccus* (N=16), were the genera with the greatest numbers of species followed by *Prochlorococcus* (N=4), *Riococcus* (N=3) and *Thaumococcus* (N=2) (Figure 2, Table 1 and S1). The AAI among PC members varied considerably, between 57 and 99.9%. The closest genus to *Prochlorococcus marinus* CCMP1375^T^ was *Riococcus* MIT9211^T^ having 69.4% AAI. The percentages of AAI variability between genera and species were 57 to 69.9% and 70 to 94%, respectively. The percentages of AAI variability within genera and species were >70 to 94% and 95 to 99.9%, respectively.

**Figure 2.**
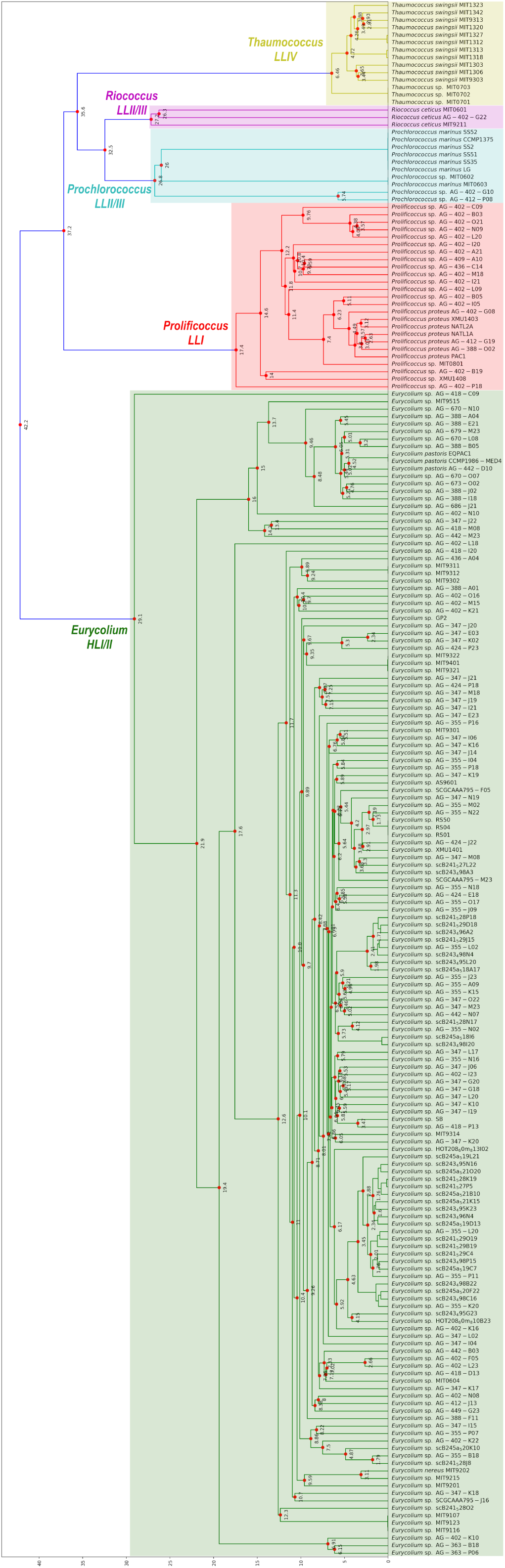
Hierarchical clustering based on average amino acid identity (AAI) analysis of the 208 PC genomes. The intraspecies limit is assumed as ≥95%, whereas genera delimitation is assumed as ≥70%. The colors (green, red, blue, purple and yellow) represent the five genera.

### Phylogenomic framework for the PC

To establish the phylogenetic structure of the five PC genera, phylogenetic analysis was performed using a four-locus based tree (MLSA) and 249 core protein sequences. Bootstrap analysis indicated that most of the branches were highly significant. These analyses have a much higher resolution than the 16S rRNA phylogeny. The PC harbours very similar 16S rRNA gene sequences with similarity ranging from 97 to 99% [2]. The MLSA and the core protein tree showed the relationship of the five genera (Figure 3). The MLSA and the core protein sequence identities among PC members varied considerably, from 49 to 100% and from 56 to 100%, respectively. The type species of the genus *Prochlorococcus, P. marinus* CCMP1375^T^, showed identity maxima of 80% and 70% MLSA and core protein sequence similarity, respectively with all other genera. The variability between species was < 95, while that within species ranged from 96 to 100% similarity. The phylogenetic analysis based on MLSA and core protein sequences (Figure 3) demonstrated congruence between the genetic and ecological features of the five novel genera. For instance, *Eurycolium* comprises high-light (HLI/II/VI) ecotypes with 30-33% GC contents, whereas the genera *Prolificoccus* and *Prochlorococcus* comprise low-light (LLI and LLII/III, respectively) ecotypes with 34-35% and 36-37% GC contents, respectively. *Riococcus* and *Thaumococcus* consists of low-light ecotypes (LLII/III and LLIV, respectively) with 37 to 38% and 50 to 50.7% GC contents, respectively.

**Figure 3.**
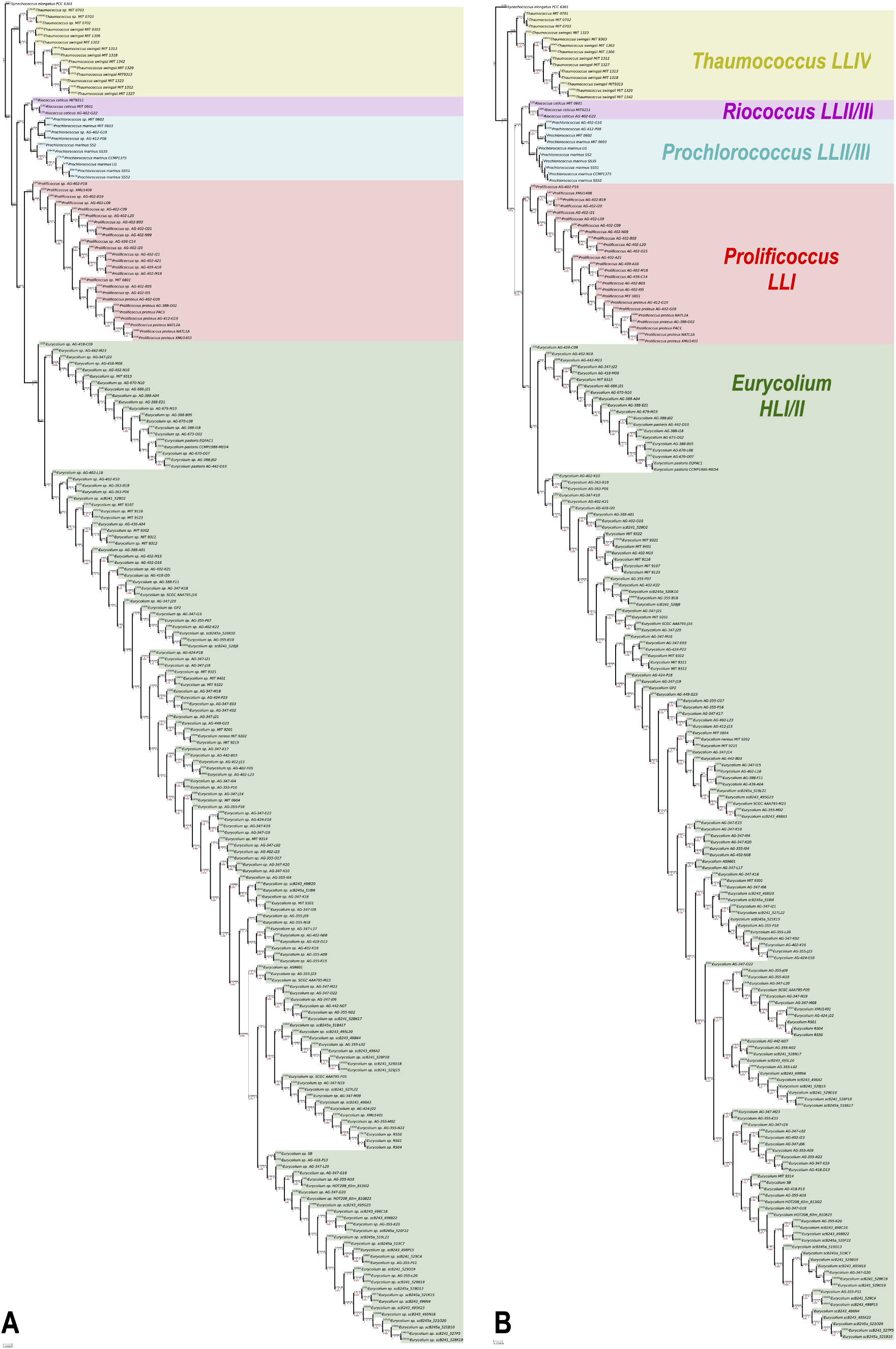
Phylogenetic trees using (A) 249 core protein sequences and (B) four house-keeping genes (*gyrB, pyrH, recA, rpoB*) based on Maximum Likelihood method. Bootstrap test was conducted using 1,000 replicates. The tree outgroup is *Synechococcus elongatus* PCC 6301.

### *In silico* phenotype prediction

Useful phenotypic features that may be used for the identification and classification of the PC were identified, including nitrogen, phosphorous, arsenate, and iron utilization (Table 2). The genus *Prochlorococcus* does not present some nitrogen-related genes (*narB, nirA, cynS, ureA*, and *gdhA*) and phosphorous-related genes (*phoB, ptrA and phoR*) but some genomes do present other nitrogen-related genes (*ntcA, glnB, amt, glnA, glsF*), an arsenate-related gene (*arsC*) and iron-related genes (*piuC* and *som*). *Prochlorococcus* is differentiated from its closest genus *Riococcus* by the presence of the *arsC* gene. All genera present the gene *Pro1404* that confers the capability for glucose uptake in PC [60].

**Table 2.** Predicted phenotype of new proposed genera of PC. “+” Presence, “-” absent, “V” variable. Nitrogen-related genes: narB, nirA, ntcA, glnB, amt, cynS, ureA, glnA, glsF, gdhA. Phosphorous-related genes: phoB, ptrA, phoR. Arsenate-related gene: arsC. Iron-related genes: piuC, som. a: Data extracted from Berube et al (2018,2019).

| Genus | Ecotype<sup>a</sup> | *narB* | *nirA* | *ntcA* | *glnB* | *amt* | *cynS* | *ureA* | *glnA* | *glsF* | *gdhA* | *phoB* | *ptrA* | *phoR* | *arsC* | *piuC* | *som* |
| --- | --- | --- | --- | --- | --- | --- | --- | --- | --- | --- | --- | --- | --- | --- | --- | --- | --- |
| *Eurycolium* | HLI/II/VI | v | v | + | v | + | v | v | + | + | v | v | - | - | v | - | + |
| *Prochlorococcus* | LLII/III | - | - | v | v | v | - | - | v | v | - | - | - | - | v | v | v |
| *Prolificoccus* | LLI | v | v | v | v | v | v | v | v | v | - | - | - | - | v | v | v |
| *Thaumococcus* | LLIV | - | v | v | v | v | - | v | v | v | v | - | v | v | v | v | v |
| *Riococcus* | LLII/III | - | - | v | v | v | - | - | v | v | - | - | - | - | - | v | v |

Overall, we demonstrated that the methods used (AAI, core protein sequences and MLSA) correlated and provided significant taxonomic resolution for differentiation of genera and species in PC (Figure 2 and 3). The proposed genera are in complete agreement with previous literature and support previous theories on the evolution of Prochlorococcus [11, 13, 14, 19, 45, 47, 50]. The PC genomic signatures were more similar between closely related species than distantly related species. The analysis performed in this study establishes a new taxonomic framework for PC.

### Challenges in the taxonomy of the Cyanobacteria phylum and the PC

Through evolution, Cyanobacteria became one of the most diverse and widely distributed prokaryote groups, occupying many niches within terrestrial, planktonic, and benthic habitats. Its long history has resulted in the evolution of broad heterogeneity, comprising unicellular and multicellular, photosynthetic and non-photosynthetic, free-living, symbiotic, toxic and predatory organisms [110–113]. Cyanobacteria have been named according to the Botanical Code [114]. The inclusion of Cyanobacteria in taxonomic schemes of Bacteria was only proposed in 1978 by Stanier et al [115]; over time, bacterial taxonomic names have come into conflict with the botanical nomenclature [116, 117]. More than two decades passed before the Note to General Consideration 5 (1999) was published indicating that Cyanobacteria should be included under the rules of the International Committee on Systematic Bacteriology (ICSB)/International Committee on Systematic of Prokaryotes (ICSP) [118–120]. Taxon nomenclature within this group has long been a topic of discussion, but there is currently no consensus [121–124]. As a result, more than 50 genera of Cyanobacteria have been described since 2000, and many of them remain unrecognized in the List of Prokaryotic Names with Standing in Nomenclature (LPSN, http://www.bacterio.net) [125] and in databases (e.g., NCBI) or have been the target of high criticism [126, 127].

Cyanobacterial taxonomy is based on morphologic traits and may not reflect the results of phylogenetic analyses [110, 128–132]. There is a predominance unrelated Cyanobacteria morphologically assembled into polyphyletic species, genera and higher taxonomic categories, which will require revisions in the future [133]. The polyphyly is indicative of the taxonomic mislabelling of many taxa. The analysis of 16S rRNA gene sequences is useful for charting and characterizing microbial communities [134] but lacks sensitivity to evolutionary changes that occur in association with ecological dynamics, in which microbial diversity is organized by physicochemical parameters [135, 136]. Hence, the processes that shape cyanobacterial communities over space and time are less well known. A recent study proposed that there should be 170 genera of Cyanobacteria based only on 16S rRNA sequences [134]. Farrant et al. delineated 121 ecologically significant taxonomic units (ESTUs) of *Prochlorococcus* and 15 ESTUs *Synechococcus* in the global ocean using single-copy *petB* sequences (encoding cytochrome b6) and environmental cues [137]. Although it was assumed that all the disclosed *Prochlorococcus* belonged to a single genus, it was not clear how many phylogenetic groups (distinct genera) corresponded to these 121 *Prochlorococcus*. Our study suggests that PC comprises at least five genera and at least 122 species. The 121 ESTUS obtained correspond to our five genera and 121species.

### Unlocking the genomic taxonomy of the PC

Despite the astonishing advances in understanding the ecology of the PC, its taxonomic structure has remained puzzling until recently. The *Prochlorococcus* collective is thought to present a high degree of panmixis due to horizontal gene transfer and is composed of one genus and one species comprising two subspecies, *P. marinus* subsp *marinus* and *P. marinus* subsp *pastoris* [47, 138–141]. We remark that at least 12 stable ecotypes have been delineated in ecological studies. On the other hand, the taxonomic heterogeneity among the named HL and LL ecotypes was noted by Thompson et al., and the current ecotypes were split into 10 species based on species definition as a group of strains that share >95% DNA identity in MLSA, >95% AAI, >70% identity of GGD and a maximum GC% range 2%. Strains of the same species form monophyletic groups on the basis of MLSA [16]. Walter et al. [18] further split the *Prochlorococcus* collective into three new genera: (1) the genus *Eurycolium* (type species *Eurycolium pastoris* with the type strain MED4^T^ (= CCMP 1986^T^), *E. chisholmii* with its type strain AS9601^T^, *E. ponticus* with its type strain MIT9301^T^, *E. nereus* with its type strain MIT9202^T^), *E. neptunius* with its type strain MIT9312^T^, *E. ponticus* with its type strain MIT 9301^T^, and *E. tetisii* with its type strain MIT9515^T^; (2) the genus *Prolificoccus*, with *Prolificoccus proteus* as the type species and NATL2A^T^ as its type strain; and (3) the genus *Thaumococcus*, with *Thaumococcus swingsii* as the type species and the type strain MIT9313^T^. The current (type) genus *Prochlorococcus* contains two species: the type species *Prochlorococcus marinus* and its type strain SS120^T^ (= CCMP 1375^T^) and *P. ceticus* with its type strain MIT9211^T^. The genus *Eurycolium* seems to form a large diverse eco-genomic group related to high temperature and oligotrophic environments, whereas the genera *Prochlorococcus* and *Prolificoccus* seem to be abundant at low temperature; *Thaumococcus* appears to thrive at low temperature and in copiotrophic environments [18].

On the basis of the eco-genomic and in silico phenotype evidence gathered amassed here, we propose five new genera within the PC. The taxonomic structure of the PC now comprises the following type species and type strains or type sequences: (1) the genus *Prochlorococcus* (Pro’chlo ro coc”cus. Gr. pref. pro, before (primitive); Gr. adj chloros, green; M.L. masc n. coccus, berry. Primitive green sphere) (the type genus). The type species *Prochlorococcus marinus* (ma’ri”nus. marinus L. adj., marine) with its type strain CCMP1375^T^ (=SS120^T^). The genome of this strain presents a GC content of 36,4%; (2) the genus *Prolificoccus* (Pro.li.fi.co.ccus. L. prolificus, productive, abundant, numerous; *Prolificoccus*, referring to an abundant coccus). The type species *Prolificoccus proteus* (L. n. proteus, ofprotos, “first”, is an early sea god, the old man of the sea) with its type strain NATL2A^T^. The genome of this strain presents a GC content of 35,1%; (3) the genus *Eurycolium* (Eur.y.co.lium. Gr. adj. eury, wide, broad; L. cole, inhabit; Gr. ium, quality or relationship, *Eurycolium*, referring to the spread inhabiting trait in marine habitats). The types species *Eurycolium pastoris* (pa.sto’ris. Latinized form of Pasteur) with its type strain CCMP1986^T^ (=MED4^T^). The genome of this strain presents a GC content of 30,8%; (4) the genus *Thaumococcus* (Thau.mo.co.ccus. Referring to Thaumas, Greek god of the wonders of the sea; N.L. masc. n. coccus (from Gr. Masc. n. kokkos, grain, seed, kernel); N.L masc. n. *Thaumococcus*, referring to a coccus living in the sea). The type species *Thaumococcus swingsii* (L. gen. masc. n. *swingsii*, of Swings, in honor of the Belgian microbiologist Jean Swings) with its type strain MIT 9313^T^. The genome of this strain presents a GC content of 50,7%; and (5) the genus *Riococcus* (Rio.coc’cus. N.L. masc. n. Rio, of Rio de Janeiro, a city in Brazil; N.L. masc. n. kokkos, a grain; N.L. masc. n. *Riococcus*, referring the city of Rio de Janeiro). The type species *Riococcus ceticus (*L. n. *ceticus*, of Cetus, denotes a large fish) with its type strain MIT9211^T^. The genome of this strain presents a GC content of 38%; We suggest that a PC species may be defined as a group of strains that share >95% DNA identity in core protein sequences, and >95% AAI. Strains of the same species will form monophyletic groups on the basis of core protein sequences.

## CONCLUSIONS

The PC represents an excellent model for further theory development in ecology, evolution, phylogeny and taxonomy. The PC refers to a group of picocyanobacteria that radiated from *Synechococcus* during millions of years of evolution in the oceans. The taxonogenomic analysis of 208 genomes revealed at least 5 new genera in addition to the original genus, *Prochlorococcus* [142]. As we delineated genera only for those groups with at least three available genome sequences, we anticipate that new genera will be included in our new proposal when more representative genomes are added to groups with only 1 or 2 genomes. On average, 20% of the genes of PC genomes have unknown functions and await further characterization. The pan-genome of the PC is estimated to include over 80000 genes [14, 46].

Because of the universal occurrence of the PC in the oceans, it may serve as a basis for monitoring the health of the oceans and climate change. Ecological transitions and changes in the oceans trigger natural genetic engineering and genome changes on a massive scale [25, 143]. The PC has evolved in multiple ecologic niches in the global ocean and participates in relevant biogeochemical cycles, such as the recently postulated parasitic arsenic cycle between *Eurycolium* and *Pelagibacter ubique* [144, 145]. We demonstrated here that the current PC is a diverse group of genera that are clearly distinguishable by eco-genomics. Ecological studies on the PC may need to consider the diversity of genera and species when novel ecologic and evolutionary hypotheses are tested.

## ACKNOWLEDGMENTS

The authors thank CNPq, CAPES and FAPERJ.

## Notes

#### Summary of Updates

The original version had a typo in the title, it has now been fixed.

